# Multiomics Analysis of the mdx/mTR Mouse Model of Duchenne Muscular Dystrophy

**DOI:** 10.1101/589424

**Authors:** Douglas W Van Pelt, Yalda A Kharaz, Dylan C Sarver, Logan R Eckhardt, Justin T Dzierzawski, Nathaniel P Disser, Alex N Piacentini, Eithne Comerford, Brian McDonagh, Christopher L Mendias

## Abstract

Duchenne muscular dystrophy (DMD) is a progressive neuromuscular disease characterized by extensive muscle weakness. Patients with DMD lack a functional dystrophin protein, which transmits force and organizes the cytoskeleton of skeletal muscle. Multiomic studies evaluate combined changes in the transcriptome, proteome, and metabolome, and have been proposed as a way to obtain novel insight about disease processes from preclinical models. We therefore sought to use this approach to study pathological changes in dystrophic muscles. We evaluated hindlimb muscles of male mdx/mTR mice, which lack a functional dystrophin protein and have deficits in satellite cell abundance and proliferative capacity. Wild type (WT) C57BL/6J mice served as controls. Muscle fiber contractility was measured, along with changes in the transcriptome using RNA sequencing, and in the proteome, metabolome, and lipidome using mass spectroscopy. While mdx/mTR mice displayed gross pathological changes and continued cycles of degeneration and regeneration, we found no differences in fiber contractility between strains. However, there were numerous changes in the transcriptome and proteome related to protein balance, contractile elements, extracellular matrix, and metabolism. There was only a 53% agreement in fold change data between the proteome and transcriptome, highlighting the need to study protein abundance along with gene expression measures. Numerous changes in markers of skeletal muscle metabolism were observed, with dystrophic muscles exhibiting elevated glycolytic metabolites. These findings highlight the utility of multiomics in studying muscle disease, and provide additional insight into the pathological changes in dystrophic muscles that might help to guide evidence-based exercise prescription in DMD patients.

## Introduction

Duchenne muscular dystrophy (DMD) is a rapidly progressing neuromuscular disease characterized by profound muscle weakness that leads to a near total loss of mobility, as well as other comorbidities that severely limit quality of life (20, 38). The dystrophin gene, which is defective in patients with DMD, encodes a protein that plays a central role in transmitting forces generated by myofibrils within muscle fibers out to the extracellular matrix (ECM) which surrounds the muscle cell. The lack of dystrophin causes membrane shearing and other structural damage to muscle cells, which then leads to inflammation, scar tissue formation, and an eventual loss of muscle fibers (7, 17, 36). Although there have been numerous innovations in the treatment of patients with DMD that have provided some improvements in muscle function and extended lifespan, the additional years gained in lifespan are often spent in a tetraplegic state in which patients are entirely dependent upon others for their care (20, 38). Therefore, there is substantial room for improvement in our ability to treat the loss of strength in individuals with DMD. Therapeutic exercise is an emerging therapy in the treatment of patients with DMD, with the goal of extending the time patients have before ambulation is lost (26). Understanding the metabolic and force generating capacity of muscle tissues can help to guide the evidence-based design of exercise programs, and may have utility in improving the treatment of patients with DMD (21).

Mouse models of disease can serve an important role in providing additional information about the etiology of disease processes and can also be useful to screen potential therapies prior to conducting clinical trials. The mdx mouse contains a premature stop codon in exon 23 of the dystrophin gene resulting in a loss of a functional dystrophin protein, and this strain is the most commonly used preclinical model for the study of DMD (34). While mdx mice lack dystrophin, they have a generally mild limb muscle phenotype that does not mimic the pathological changes observed in patients with DMD (34). This deficiency has limited the utility of this model, and led to the development of additional mouse models with a phenotype that more closely parallels the human disease (34). One reason that mdx mice do not show the progressive loss in muscle quality that is observed in patients with DMD is due to species-specific differences in satellite cells, which are the myogenic stem cells responsible for regenerating injured fibers (16). Although dystrophin does play a role in self-renewal and proper asymmetric cell division (10), murine but not human satellite cells pools are thought to avoid depletion by inducing the expression of telomerase genes which restore telomere length that is severely shortened in response to proliferating throughout continued cycles of degeneration and regeneration (39). To address this and improve the mdx model, a new line of mice was generated in the mdx background in which the telomerase RNA component (Terc) gene was deleted (39). These genetically modified animals, referred to as mdx/mTR mice, display reduced satellite cell abundance and proliferative capacity, and a more pronounced muscle degenerative phenotype than mdx mice (39).

Multiomic studies, which comprehensively evaluate the transcriptome, proteome, and metabolome, have been proposed as a way to gain greater translational insight from preclinical models of human disease (2). While numerous studies have evaluated isolated histological, biochemical, and contractile changes in the muscles of animal models of muscular dystrophy, and some limited individual proteomic, metabolomic, or transcriptomic studies have been performed, there is a general lack of connecting physiological measures with multiomic techniques. Therefore, our goal was to perform an integrated and comprehensive analysis of muscle cell contractility and histological features with the transcriptome, proteome, metabolome, and lipidome of 4-month old male mdx/mTR and wild type (WT) C57BL/6J mice.

## Methods

### Animals

This study was approved by the University of Michigan IACUC (protocol PRO00006079), and all experiments were performed in accordance with the US Public Health Service (PHS) Policy on Humane Care and Use of Laboratory Animals. Mice were obtained from Jackson Labs (Bar Harbor, ME, USA) and housed under specific pathogen free conditions. Mice were allowed *ad libidum* access to food and water, but were fasted for four hours prior to harvest to minimize any impacts of acute food consumption on metabolomic measurements. We utilized male mdx/mTR^G2^ mice (referred to as mdx/mTR mice in this paper, Jackson Labs strain 023535) which are deficient for both the dystrophin and telomerase RNA component (Terc) genes (39). Wild type male C57BL/6J mice (referred to as WT mice in this paper, Jackson Labs strain 000664), which is the same background of the mdx/mTR mice, were used as controls.

DNA obtained from tail biopsies, and genotype of animals was verified by PCR analysis of following guidelines of Jackson Labs. Mice were obtained at two months of age and were closely monitored for gross cage activity and responsiveness to touch stimuli. As mice approached four months of age, there was a marked decline in their apparent mobility and responsiveness to stimuli. We therefore selected four months of age for analysis, as we suspected this would likely be reflective of the critical time point that patients with DMD transition from ambulatory to non-ambulatory status, and therefore a translationally relevant time point for dystrophic pathology.

At four months of age, mice were injected with sodium pentobarbital via an intraperitoneal route to obtain deep anesthesia, tissues were harvested and weighed, and animals were humanely euthanized by anesthetic overdose followed by cervical dislocation. The tissues collected from both hindlimbs included the extensor digitorum longus (EDL), used for single fiber testing and histology; gastrocnemius (gastroc) muscles, used for histology; plantaris muscles, used for proteomics; soleus muscles, used for histology; and tibialis anterior (TA) muscles used for metabolomics/lipidomics and RNAseq/gene expression. Several muscles were analyzed per animal to be consistent with the PHS guidelines on reducing the numbers of animals used in studies by maximizing the tissue sampled per animal. A total of N=6 mice were used from each genotype, for a total of N=12 mice in the study.

### RNA Sequencing (RNAseq) and Gene Expression

RNAseq and gene expression was performed as described (14). RNA was isolated from TA muscles using a miRNEasy kit (Qiagen, Valencia, CA, USA). RNA quality was assessed using a TapeStation (Agilent, Santa Clara, CA, USA) and all samples had an RNA integrity number greater than 9.0. For each sample, 250ng total RNA was delivered to the University of Michigan Sequencing Core for mRNA sequencing (RNAseq) analysis. Sample concentrations were normalized, cDNA pools were created for each sample and cDNAs were then subsequently tagged with a barcoded oligo adapter to allow for sample specific resolution. Sequencing was carried out using an Illumina HiSeq 4000 platform (Illumina, San Diego, CA, USA) with 50bp single end reads and an average of approximately 65 million reads per sample. Quality assessment and fold change calculations were performed in BaseSpace (Illumina). Sequencing data was quality checked using FastQC, aligned to the reference genome (mm10, UCSC, Santa Cruz, CA, USA) with the STAR software package, and differential expression based on fragments per kilobase of transcript per million mapped reads was performed using DESeq2. Genes with a base mean expression of at least 10 were used in analysis. A total of 14064 transcripts met this threshold. RNAseq data has been deposited to NIH GEO (accession GSE127929).

For gene expression analysis, RNA was reversed transcribed into cDNA using iScript Reverse Transcription Supermix (Bio-Rad). Quantitative PCR (qPCR) was performed with cDNA in a CFX96 real-time thermal cycler (Bio-Rad) using iTaq Universal SYBR Green Supermix (Bio-Rad). Target gene expression was normalized to the housekeeping gene β2-microglobulin, and further normalized to WT samples using the 2^-ΔΔCt^ method.

### Proteomics

Proteomics was performed as reported in previous studies (3, 27, 40) at the University of Liverpool Centre of Genomic Research. Protein extraction of plantaris muscles was performed by first extracting soluble proteins from the homogenization supernatant, followed by second protein extraction of the remaining insoluble pellet of the muscle. Soluble proteins were extracted by homogenizing tissue in 25mM ammonium bicarbonate and 10mM iodoacetamide, followed by centrifugation and removal of the supernatant. Then 100µg of soluble protein was further reduced and alkylated, and in solution trypsin digestion was performed. The remaining pellet was resuspended in 500µl of 4M Guanidine-HCl extraction buffer (GnHCl), 65mM dithiothreitol, 50mM sodium acetate, pH 5.8 for 48 h at 4°C with shaking. The samples were then centrifuged, the supernatant was removed, and 50µg of the soluble GnHCl fraction was subjected to in solution-trypsin digest on 10µl of resin (StrataClean, Agilent, Cheshire, UK) followed by reduction and alkylation. The digests from the isolated soluble proteins (5µL, corresponding to 2.5µg of peptides) and of those from GnHCl fraction (10µL, corresponding to 5 µg peptides) were loaded onto a spectrometer (Q-Exactive Quadrupole-Orbitrap, Thermo Scientific, Loughborough, UK) on a one-hour gradient with an intersample 30-minute blank loaded.

Raw spectra were converted to mgf files using Proteome Discovery (Thermo Scientific, Loughborough, UK) and resulting files were searched against the UniProt mouse sequence database using a Mascot server (Matrix Science, London, UK). Search parameters used were: peptide mass tolerance, 10ppm; fragment mass tolerance, 0.01Da; +1,+2,+3 ions; missed cleavages 1; instrument type ESI-TRAP. Variable modifications included carbamidomethylation of cysteine residues and oxidation of methionine. Label free quantification was performed using PEAKS Studio software (BSI, Waterloo, ON, Canada). Proteins identified from the soluble and the insoluble fraction were combined for each group, and MetaboAnalyst 4.0 (6) was used to quantify differences between groups. The full proteomics data for this study has been deposited in the ProteomeXchange Consortium via the PRIDE partner repository (accession PXD010507).

### Pathway Enrichment

The Canonical Pathway feature of Ingenuity Pathway Analysis software (Qiagen) was used to perform pathway enrichment analysis for RNAseq and proteomics data as described (9). Activation state was determined by the value of the z-score for each pathway. Pathways relevant to dystrophic pathology were selected for reporting, with the full data set available in the Supplemental Material.

### Whole Muscle Histology

Histology was performed as described (14). EDL, gastroc, and soleus muscles were snap frozen in tragacanth gum and stored at −80°C until use. Muscles were sectioned at a thickness of 10µm, fixed with 4% paraformaldehyde, and then incubated in 0.2% Triton-X 100. Sections were then stained with wheat germ agglutinin (WGA) conjugated to AlexaFluor 555 (AF555, Thermo Fisher, Waltham, MA, USA) to label the extracellular matrix, and DAPI (Sigma, Saint Louis, MO, USA) to identify nuclei. Sections were also stained with hematoxylin and eosin (H&E, Vector Labs, Burlingame, CA, USA). Images of stained sections were taken on a microscope (BX51, Olympus, Waltham, MA, USA), and ImageJ (NIH, Bethesda, MD, USA) was used to quantify fiber CSAs and the percentage of muscle fibers with centrally located nuclei from WGA-AF555 and DAPI images.

### Muscle Fiber Contractility

The contractility of chemically permeabilized muscle fibers was performed as described (28, 29). Briefly, fiber bundles were dissected from the EDL muscle, placed in skinning solution for 30 min and then in storage solution for 16 h at 4°C, followed by storage at −80°C. On the day of contractility assessment, samples were thawed slowly on ice, and individual fibres were pulled from bundles using fine mirror-finished forceps. Fibers were then placed in a chamber containing relaxing solution and secured at one end to a servomotor (Aurora Scientific, Aurora, ON, Canada) and the other end to a force transducer (Aurora Scientific) using two ties of 10-0 monofilament nylon suture (Ashaway Line & Twine, Ashaway, RI, USA). A laser diffraction measurement system was used to adjust fiber length to obtain a sarcomere length of 2.5µm. Mean fiber CSA was calculated assuming an elliptical cross-section, with diameters obtained at five positions along the fiber from high-magnification images taken from top and side views. Maximum isometric force (F_o_) was elicited by immersing the fiber in a high calcium activation solution. Specific maximum isometric force (sF_o_) was calculated by dividing F_o_ by fiber CSA. The susceptibility of fibers to a lengthening contraction-induced injury was assessed by applying a single stretch to fully-activated fibers. The stretch was equivalent in amplitude to 30% of fiber length, and was applied at a constant velocity of 0.5 fiber lengths per second. After the stretch, fibers were immediately returned to their original length and allowed to generate force until a new steady-state level was reached. The difference between the pre- and post-stretch forces was used to calculate force deficit, expressed as a percentage of pre-stretch F_o_. Eight to ten fast fibers were tested from each EDL muscle.

### Muscle Fiber Western Blot

Western blots of permeabilized muscle fibers was performed to evaluate the presence of dystrophin in fibers from samples that were pulled from bundles of WT and mdx/mTR muscles, as described above. Approximately 30 pulled fibers from each sample were homogenized in Laemmli’s sample buffer (Bio-Rad, Hercules, CA, USA), boiled for 2 minutes, and loaded into 7.5% polyacrylamide gels (Bio-Rad), and subjected to electrophoretic separation. Proteins were transferred to 0.2µm nitrocellulose membranes (Bio-Rad) using a semi-dry transfer apparatus (Trans-Blot Turbo, Bio-Rad). Membranes were then blocked with 5% powdered milk, and incubated with polyclonal rabbit antibodies against dystrophin (ab15277, AbCam, Cambridge, MA, USA), and goat anti-rabbit antibodies conjugated to horse radish peroxidase (A16124, Thermo Fisher). Proteins were detected using enhanced chemiluminescence reagents (Clarity, Bio-Rad) and imaged in a chemiluminescent system (ChemiDoc MP, Bio-Rad). Following visualization of proteins, membranes were briefly stained with Coomassie (Bio-Rad) to verify similar protein loading between lanes.

### Muscle Fiber Histology

To determine whether dystrophin persisted at the sarcolemma after permeabilization, fibers from WT and mdx/mTR samples were pulled from bundles that were prepared as described above, and subjected to histological processing using previously reported techniques (25). Fibers were fixed in 2% glutaraldehyde (Electron Microscopy Sciences, Hatfield, PA, USA), rinsed, and then incubated with phalloidin conjugated to AlexaFluor 555 (Thermo Fisher) to identify actin, polyclonal rabbit antibodies against dystrophin (ab15277, AbCam), and DAPI (Sigma) to label nuclei. Goat anti-rabbit antibodies conjugated to AlexaFluor 647 (A21244, Thermo Fisher) were used to detect primary antibodies. Slides were imaged using a laser scanning confocal microscope (LSM 880, Zeiss, Thornwood, NY, USA).

### Metabolomics and Lipidomics

The University of Michigan Metabolomics Core performed mass spectrometry-based shotgun lipidomics and metabolomics measurements from snap frozen, homogenized muscle samples as described (14, 41). For lipidomics, lipids were extracted from samples with a solvent mixture consisting of 2:2:2 (v/v/v) methanol:dichloromethane:water mixture at room temperature after adding internal standard mixture. Once dried, the samples were resuspended in a solution containing 1:5:85 (v/v/v) acetonitrile:water:isopropanol and 10mM ammonium acetate. Samples were then subjected to liquid chromatography-mass spectrometry (LC-MS), and MS peaks were matched in-silico with LipidBlast (18). Quantification was performed by Multiquant software (AB-SCIEX, Framingham, MA, USA). One WT sample failed quality control and was excluded from analysis.

For metabolomics, metabolites were extracted from frozen muscle in a solvent mixture containing 8:1:1 methanol:chloroform:water (v/v/v). Metabolites were derivatized and analyzed with gas chromatography-MS (22). Quantification of metabolites was performed using Masshunter Quantitative Analysis software (Agilent Technologies, Santa Clara, CA, USA).

MetaboAnalyst 4.0 (6) was used to perform detailed statistical analyses for lipidomics and metabolomics assays. Metabolomics and lipidomics data has been deposited to the UCSD Metabolomics Workbench (study ST000753).

### Statistics

Values are presented as mean±SD, with the exception of qPCR which is mean±CV. Differences between groups were assessed using t-tests (α=0.05). For RNAseq, proteomics, lipidomics, and metabolomics data, a FDR-correction was applied to adjust for multiple observations. A binomial test (α=0.05) was used to compare the distribution of matched proteins and transcripts that have the same direction of fold change in mdx/mTR mice normalized to controls and in those which the direction of the fold change of the protein is different from the transcript. A Wilson/Brown approach was used to generate 95% confidence intervals for the percent distribution. A Chi-square test was used to evaluate differences between the size distribution of muscle fibers. With the exception of analyses performed in DESeq2 or MetaboAnalyst, statistical calculations were performed using Prism software (version 8.0, GraphPad, San Diego, CA, USA).

### Supplemental Data

A data supplement containing processed RNAseq, proteomics, metabolomics, lipidomics, IPA analyses, and qPCR primer sequences is available as Supplemental Tables S1-S6.

## Results and Discussion

### Overview

We first sought to evaluate pathological changes in the hindlimb muscles of 4-month old WT and mdx/mTR mice, as the function of these muscles is important in maintaining ambulation. The muscles that were analyzed include the extensor digitorum longus (EDL), gastrocnemius (gastroc), plantaris, tibialis anterior (TA) and soleus. The EDL, gastroc, plantaris, and TA are composed of mostly fast (type II) muscle fibers, while the soleus has a mixed distribution of slow (type I) and fast fibers (4). Although we used different muscles to minimize the number of animals required to conduct this study, there is a substantial amount of similarity in the transcriptome of EDL, gastroc, plantaris, and TA muscles (44), and we think that conclusions from one of these fast fibered muscle groups can be extended to other muscles with similar fiber type distributions.

Translation initiation and elongation are complicated processes in skeletal muscle tissue, and the pathways that regulate protein synthesis can be distinct from those regulating gene expression (12). Therefore, to gain a comprehensive view of changes in the transcriptome and proteome of skeletal muscles of mdx/mTR mice compared to WT mice, we performed mRNA sequencing (RNAseq) using TA muscles, and untargeted mass spectrometry-based proteomics of plantaris muscles. The principal component (PC) analysis of transcriptome data demonstrate divergence between WT and mdx/mTR mice (Figure 1A). A total of 14064 transcripts were detected and met appropriate expression cut-off thresholds, with 1791 transcripts demonstrating at least a 1.5-fold increase in expression and P<0.05, and 665 transcripts were at least 1.5-fold downregulated and significantly different (Figure 1B). To further increase the number of proteins that were analyzed, we analyzed the supernatant of homogenized plantaris muscles, as well as proteins extracted from the insoluble fraction. This resulted in the detection of 12 additional proteins, in addition to 83 proteins that were already detected in the soluble fraction. Similar to the transcriptome, PC analysis of the proteome of WT and mdx/mTR groups also demonstrated divergence (Figure 2A). Of the 1237 proteins detected, 460 proteins had at least a 1.5-fold increase in abundance and P<0.05, and 25 proteins were at least 1.5-fold downregulated and significantly different (Figure 2B). We then evaluated changes in the transcriptome compared with the proteome and identified 1119 proteins that also had transcripts that were detected in RNAseq (Figure 3A). Of these 1119 protein:transcript pairs, 53.3% (95% confidence interval, 50.3-56.2%) had a fold change that was in the similar direction, while 46.7% (95% confidence interval, 43.8-49.7%) had a fold change of protein that did not match the direction of the transcript (Figure 3A). This distribution was significantly different than a random 50% distribution (Figure 3A). We then identified genes and proteins that have important roles in protein synthesis and degradation (Figure 3B), muscle fiber contractility and cytoskeletal architecture (Figure 3C), ECM (Figure 3D), and metabolism (Figure 3E), which will be discussed in subsequent sections.

**Figure 1.**
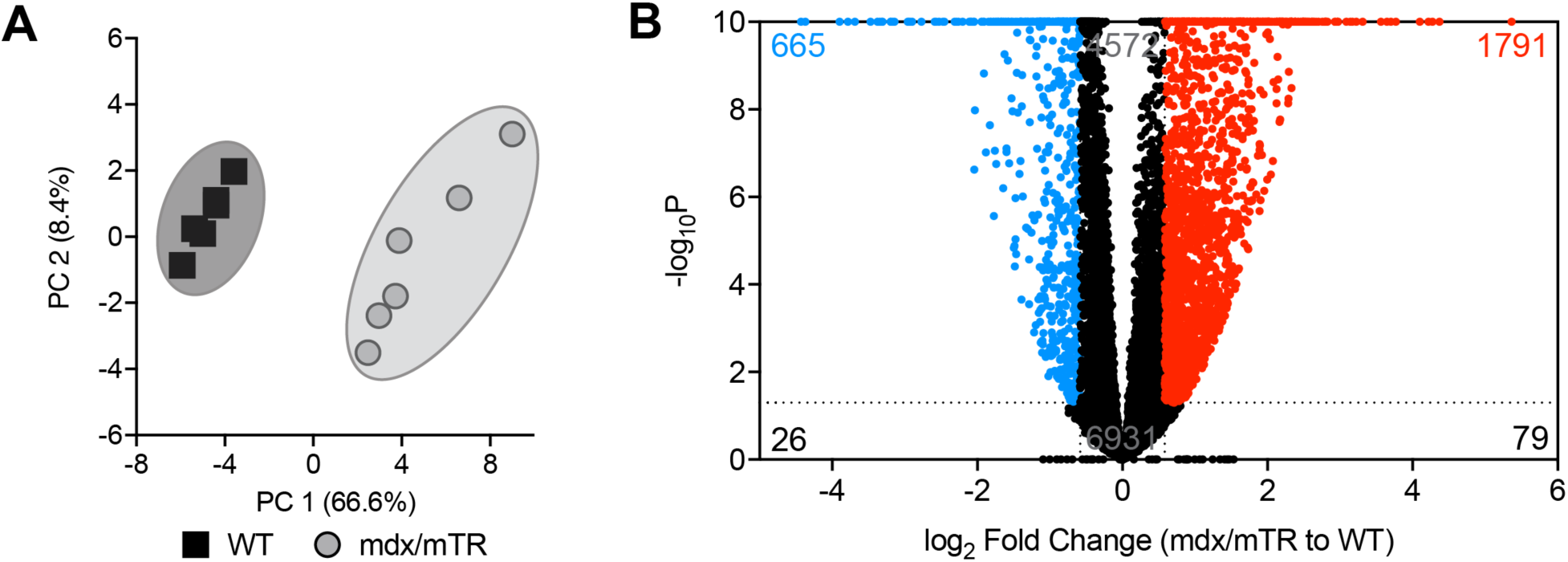
RNA sequencing. (A) Principal component (PC) analysis of RNA sequencing data. (B) Volcano plot demonstrating log_2_ fold-change and FDR-corrected P-values of all measured transcripts. Genes with -log_10_P values greater than 10 are shown directly on the top border of the graph. Genes with a > 1.5-fold upregulation in mdx/mTR mice (log_2_ fold change > 0.585) and P value < 0.05 (-log_10_P > 1.301) are shown in red. Genes with a > 1.5-fold downregulation in mdx/mTR mice (log_2_ fold change < −0.585) and P value < 0.05 (-log_10_P > 1.301) are shown in blue. N=6 WT and N=6 mdx/mTR mice per group.

**Figure 2.**
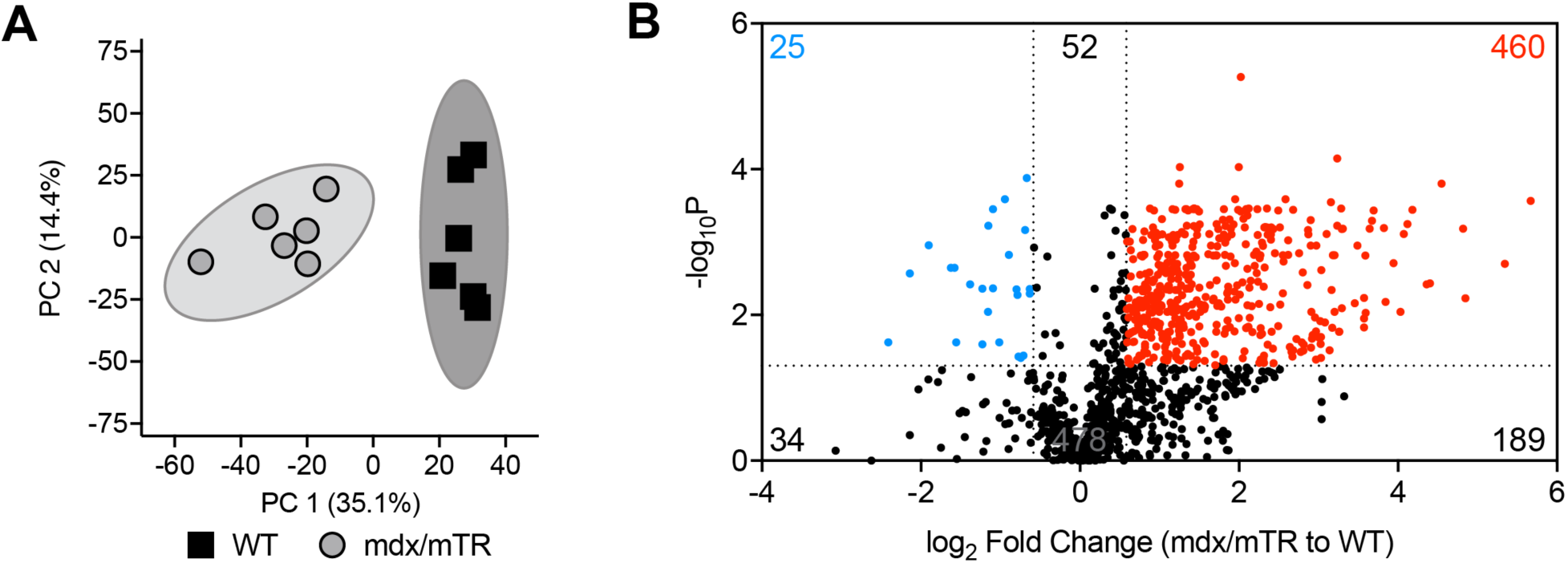
Proteomics. (A) Principal component (PC) analysis of proteomics data. (B) Volcano plot demonstrating log_2_ fold-change and FDR-corrected P-values of all measured proteins. Proteins with a > 1.5-fold increase in mdx/mTR mice (log_2_ fold change > 0.585) and P value < 0.05 (-log_10_P > 1.301) are shown in red. Genes with a > 1.5-fold decrease in mdx/mTR mice (log_2_ fold change < −0.585) and P value < 0.05 (-log_10_P > 1.301) are shown in blue. N=6 WT and N=6 mdx/mTR mice per group.

**Figure 3.**
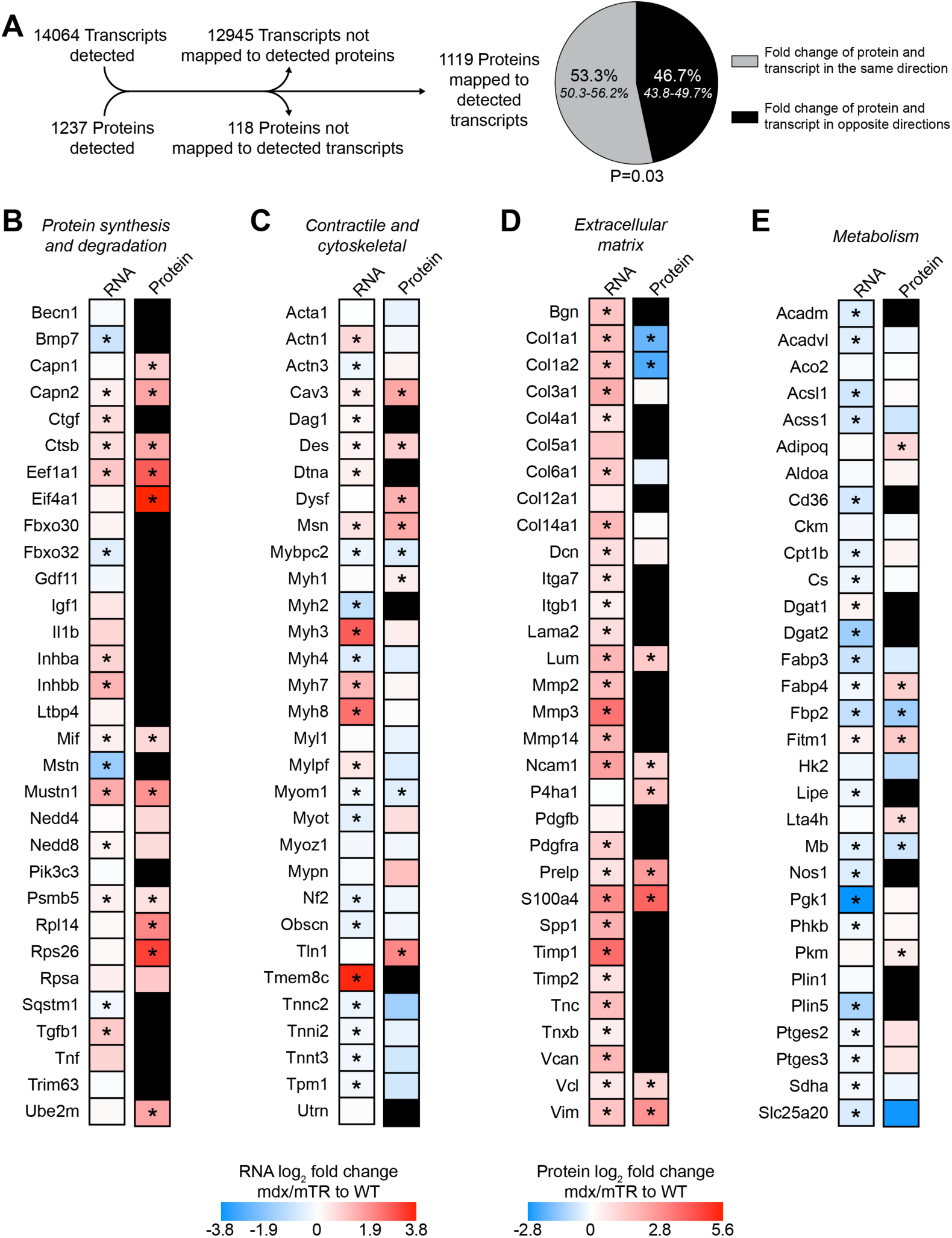
Integrated RNA sequencing and Proteomics. (A) Overview of the total number of transcripts and proteins detected, the number of proteins that matched to transcripts, and the percentage (with 95% confidence interval) of matched proteins and transcripts that have the same direction of fold change in mdx/mTR mice normalized to controls (gray), and those in which the direction of the fold change of the protein is different from the transcript (black). A binomial test was performed to compare the observed distribution to the expected distribution. Heatmap demonstrating the log_2_ fold change in selected genes and proteins related to (B) protein synthesis and degradation, (C) contractility and the cytoskeleton, (D) the extracellular matrix, and (E) metabolism. Black box indicates that the protein was not present in the proteomics data set. Differences between groups (B-E) assessed using t-tests; *, significantly different (P<0.05) from WT. N=6 WT and N=6 mdx/mTR mice.

Pathway enrichment analysis of RNAseq and proteomics data was performed (Table 1). While there was general agreement in the activation state of pathways between RNAseq and proteomics data, many of the pathways showed divergent activation states between transcriptional and proteomic analysis.

**Table 1.** Pathway enrichment analysis. Pathway enrichment analysis of RNAseq and proteomics data. Data shown are p-values and the activation state of the pathway: ↑, pathway activated; ↓, pathway inhibited; ↔, activation status could not be determined. ND, pathway not detected.

| **Pathway** | **RNAseq** | **Proteomics** |
| --- | --- | --- |
| Actin Cytoskeleton Signaling | 2.14E-09 ↔ | 8.91E-09 ↑ |
| Acute Phase Response Signaling | 6.31E-04 ↑ | 2.51E-07 ↑ |
| AMPK Signaling | 7.24E-02 ↑ | 6.03E-04 ↑ |
| Axonal Guidance Signaling | 7.94E-16 ↔ | 9.55E-05 ↔ |
| BMP signaling | 1.94E-01 ↓ | 3.31E-02 ↑ |
| Calcium Signaling | 3.16E-05 ↑ | 4.37E-04 ↑ |
| EIF2 Signaling | 1.00E+00 ↑ | 1.32E-08 ↑ |
| Ephrin Receptor Signaling | 3.24E-04 ↑ | 4.90E-03 ↑ |
| ERK/MAPK Signaling | 1.23E-01 ↑ | 1.48E-10 ↑ |
| FAK Signaling | 4.37E-03 ↔ | 9.33E-04 ↔ |
| Fatty Acid β-oxidation | 4.00E-01 ↓ | 2.88E-02 ↔ |
| FGF Signaling | 1.95E-02 ↑ | 1.04E-01 ↑ |
| Gluconeogenesis | 3.98E-02 ↔ | 1.66E-02 ↔ |
| Glycolysis | 1.12E-01 ↑ | 2.00E-03 ↑ |
| IGF1 Signaling | 1.57E-01 ↓ | 1.32E-05 ↑ |
| ILK Signaling | 5.62E-05 ↑ | 3.16E-06 ↑ |
| Inflammasome Pathway | 7.08E-06 ↑ | 3.46E-01 ↔ |
| Inhibition of Matrix Metalloproteinases | 2.34E-06 ↓ | 5.63E-01 ↔ |
| Integrin Signaling | 1.26E-02 ↑ | 1.55E-07 ↑ |
| Mitochondrial Dysfunction | 1.00E+00 ↔ | 6.92E-02 ↔ |
| mTOR Signaling | 1.00E+00 ↑ | 4.17E-09 ↑ |
| NF-κB Signaling | 4.27E-04 ↑ | 5.24E-01 ↑ |
| p70S6K Signaling | 3.89E-02 ↑ | 9.55E-05 ↑ |
| Pentose Phosphate Pathway | 1.00E+00 ↔ | 3.63E-05 ↑ |
| PI3K/AKT Signaling | 5.50E-02 ↑ | 2.29E-05 ↑ |
| Protein Kinase A Signaling | 5.62E-02 ↓ | 1.35E-06 ↑ |
| Protein Ubiquitination Pathway | 1.00E+00 ↔ | 3.98E-28 ↔ |
| RhoA Signaling | 5.01E-03 ↑ | 1.12E-05 ↑ |
| STAT3 Pathway | 5.37E-07 ↔ | 5.41E-01 ↔ |
| Triacylglycerol Biosynthesis | 1.86E-01 ↑ | ND |
| Triacylglycerol Degradation | 1.68E-01 ↓ | 2.98E-01 ↔ |
| Unfolded Protein Response | 4.90E-02 ↔ | 1.62E-04 ↔ |

Additionally, qPCR analysis of a select set of genes was performed, and we generally observed similar directions in fold change directions between qPCR and RNAseq data (Table 2).

**Table 2.**
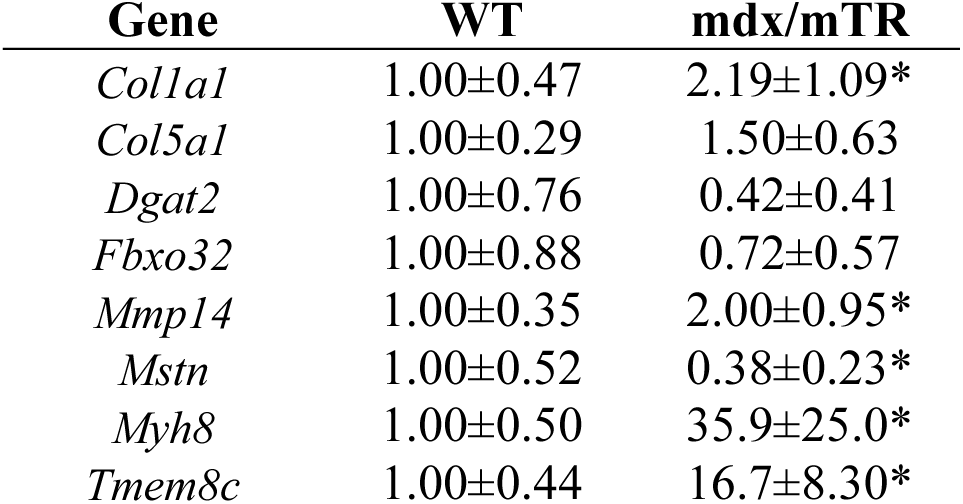
Changes in gene expression as measured by qPCR. Gene expression values from tibialis anterior muscles. Values are mean±CV, differences tested between groups using t-tests; *, significantly different (P<0.05) from WT. N=6 WT and 6 mdx/mTR mice.

### Muscle Mass, Fiber Contractility, and the Extracellular Matrix

The mass of muscles was greater in mdx/mTR mice, demonstrating an increase from 23% in EDL muscles to 45% in TA muscles (Figure 4A), which is generally similar to previous reports of mdx mice (23). This elevated muscle mass was disproportionate to the 9% higher body mass of mdx/mTR mice (26.0±1.7g for WT, 28.3±0.9g for mdx/mTR, P=0.02). While we did not observe differences between wild type and mdx/mTR mice in terms of muscle fiber cross-sectional area (CSA), the majority of muscle fibers of mdx/mTR mice contained centrally located nuclei, and fibrotic lesions were also noted along with edema between muscle fibers (Figures 4B-D). Although the mean fiber CSA was not different, the percent distribution of fibers for muscles was different (Figure 4E). These findings suggest that the increased mass of muscles of mdx/mTR mice likely occurs due to inflammation and perhaps an expansion of fibrotic ECM. This is supported by several findings from the RNAseq and proteomics data (Figure 3B-D), as well as an increase in the inflammasome pathway and MMP activity as predicted by pathway enrichment analysis of RNAseq data (Table 1).

**Figure 4.**
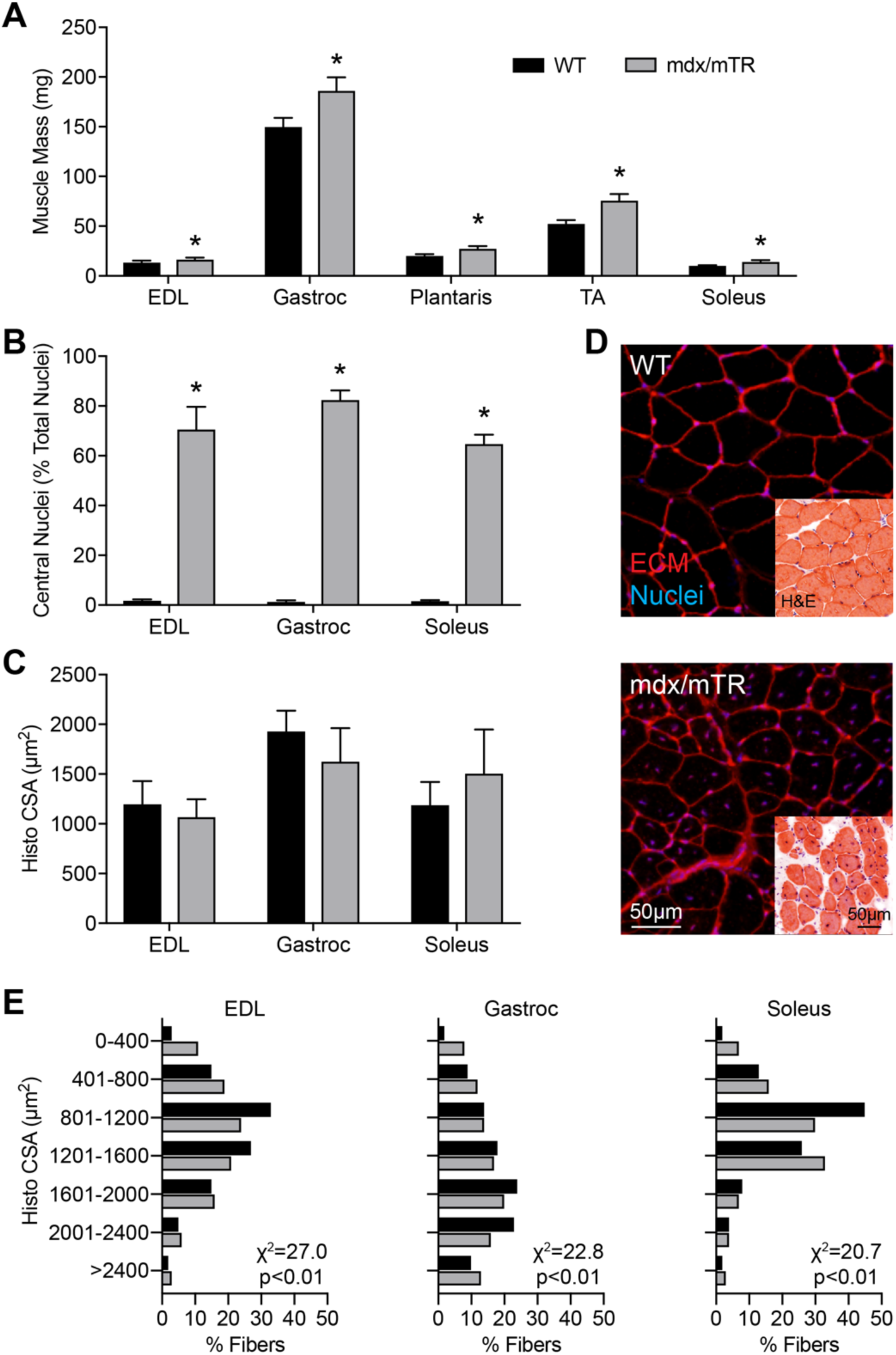
Muscle Mass and Histology. (A) Muscle mass data from extensor digitorum longus (EDL), gastrocnemius (gastroc), plantaris, tibialis anterior (TA) and soleus muscles. (B) Fibers that contain centrally located nuclei, expressed as a percentage of total fibers and (C) histological cross-sectional area (CSA). (D) Representative histology from WT and mdx/mTR gastroc muscles; red, WGA-AF555/extracellular matrix; blue, DAPI/nuclei. Inset images are muscles stained with hematoxylin and eosin (H&E). Scale bar for each respective type of image is 50µm. (E) Percent distribution of fibers by size. Values are mean+SD. Differences between groups for A-C assessed using t-tests; *, significantly different (P<0.05) from WT. Differences for E tested using a Chi-square test. N=6 WT and N=6 mdx/mTR mice.

The major skeletal muscle calpains, µ-calpain (Capn1) and m-calpain (Capn2) were elevated in mdx/mTR muscles, along with protease cathepsin B (Ctsb) (Figure 3B). Consistent with the elevation in calpains, calcium signaling was also enriched in RNAseq and proteomics data (Table 1). While the skeletal muscle specific E3 ubiquitin ligases MUSA1 (Fbxo30), atrogin-1 (Fbxo32), and MuRF-1 (Trim63) were not upregulated, the 20S proteasome subunit Psmb5 was upregulated, and signaling molecules that induce the activation of the ubiquitin ligase system, activin A (Inhba), activin B (Inhbb), and TGFβ (Tgfb1) were upregulated (Figure 3B). The protein ubiquitination pathway was not different from RNAseq data, but was among the most significantly activated pathways from proteomics data (Table 1). Several proteins or genes involved with translation were upregulated, including Eef1a1, Eif4a1, Rpl14, and Rps26 (Figure 3B). Proteomics data also demonstrated an activation of the EIF2, IGF1, mTOR, p70S6K and PI3K/Akt signaling pathways, which are all involved in the regulation of protein synthesis (16). These findings suggest that the muscles of mdx/mTR muscles are primed to synthesize new proteins to maintain muscle function, even though the cells continue to be damaged.

Based on the findings of elevated genes and proteins involved in proteolysis, and the loss in force production that is known to occur in patients with DMD (20, 38), we next sought to evaluate muscle contractility. At the whole muscle level, the specific force of intact lateral gastroc muscles from mdx/mTR mice was approximately 3.5-fold lower than WT mice (39). The lack of dystrophin protein is known to reduce whole muscle force production by disrupting force transmission from myofibrils through the dystrophin glycoprotein complex, but the specific force deficit in mdx/mTR muscles is much greater than that which is observed in mdx mice, which have a 13% reduction compared to WT mice (23). Other types of muscle injuries and chronic diseases have reduced whole muscle force production due to disruptions in myofibril force production (15, 30, 37), and therefore we measured the contractility of permeabilized muscle fibers to assess myofibril force production.

No difference in the CSAs of permeabilized fibers, maximum isometric force, or specific force was observed (Figures 5A-C). In addition to deficits in whole muscle force production, when dystrophic muscles are subjected to a lengthening contraction-induced injury in which the muscle is suddenly lengthened while maximally activated, mdx mice experience a two-fold greater force deficit than WT mice (8). Based on this observation, we performed lengthening contraction-induced injury on permeabilized muscle fibers to evaluate the damage susceptibility of myofibrils from mdx/mTR mice. No differences in force deficits after lengthening injury were observed between WT and mdx/mTR mice (Figure 5D). The findings from the current study are consistent with previous observations of contractility values of muscle fibers from mdx and WT mice (24). The lack of any appreciable difference in the susceptibility of fibers from WT and mdx mice to injury was thought to occur as a result of the permeabilization process disrupting the presence of dystrophin at the sarcolemma, although this was not directly addressed in the manuscript (24). We therefore determined whether dystrophin was properly expressed at the sarcolemma, or if it was lost during the fiber permeabilization and isolation process. Western blots for dystrophin were performed using homogenates of permeabilized fibers pulled from bundles, and the presence of dystrophin protein in WT muscle fibers was observed (Figure 5E). This was followed up by histology of permeabilized fibers demonstrating clear signal for dystrophin at the sarcolemma in WT fibers, and as expected no signal was observed for fibers from mdx/mTR mice (Figure 5F). While subsarcolemmal nuclei were present as expected in both WT and mdx/mTR mice, we also frequently observed centrally located nuclei (Figure 4B) and collections of several nuclei immediately adjacent to each other and in a central location of the fiber (Figure 5F), consistent with a recent fiber injury and myotube fusion event. Additionally, the typical myofibril striation pattern was maintained in mdx/mTR mice, suggesting that despite undergoing repeated degeneration-regeneration cycles, myofibrils are properly organized and aligned in both strains.

**Figure 5.**
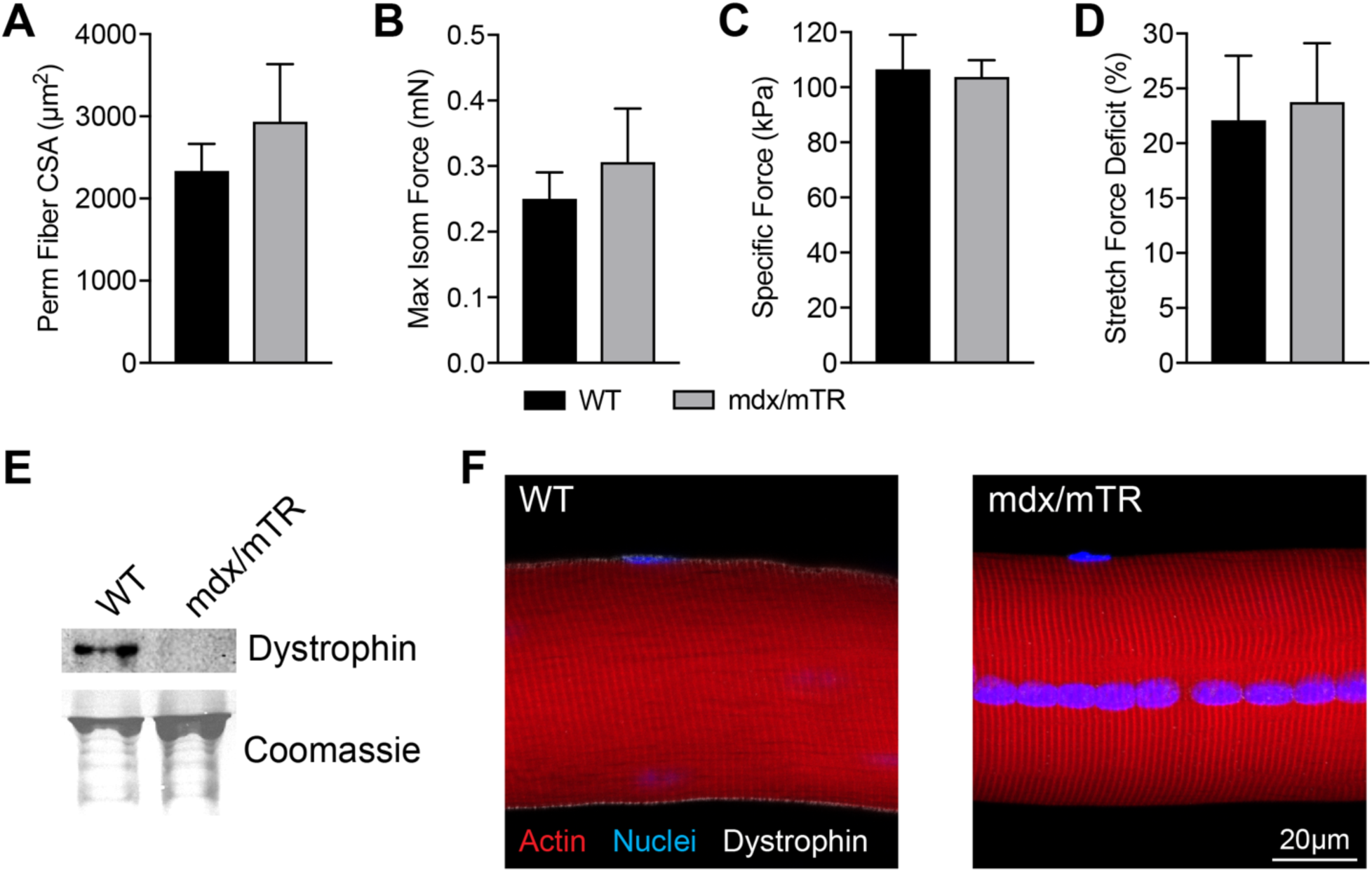
Permeabilized Extensor Digitorum Longus Muscle Fiber Force and Fiber Morphology. (A) Permeabilized fiber cross-sectional area (CSA), (B) maximum isometric force, (C) specific force, and (D) force deficit after active stretch induced injury. Values are mean+SD. Differences between groups assessed using t-tests; no differences detected between WT and mdx/mTR groups. N=6 WT and N=6 mdx/mTR mice. (E) Western blot and (F) histology of permeabilized fibers demonstrating the presence of dystrophin at the sarcolemma of fibers after the permeabilization process; red, actin; blue, DAPI/nuclei; white, dystrophin. Scale bar in both images is 20µm.

Although fiber contractile properties studies demonstrated no difference in myofibril force production, there was not a clear pattern of changes in the abundance or expression of sarcomere components such as α-actinin (Acta1), desmin (Des), moesin (Msn), myosin binding protein C (Mybpc2), myomesin (Myom1), myotilin (Myot), myozenin (Myoz1), myopalladin (Mypn), obscurin (Obscn), talin (Tln1), and the troponins (Tnn) (Figure 3C). For the myosin heavy chains, embryonic myosin heavy chain (Myh3) and perinatal myosin heavy chain (Myh8), which are expressed in regenerating fibers, were upregulated at the transcript level (Figure 3C). Pathway enrichment analysis indicated activation of cytoskeletal and mechanically regulated pathways, such as ERK/MAPK, ILK, integrin, and RhoA (Table 1). Further supporting the continued degeneration and regeneration cycle, there was an upregulation in the myoblast fusion protein myomaker (Tmem8c) and the membrane patching protein dysferlin (Dysf) (Figure 3C), which are also important in membrane repair in other models of DMD (31, 45).

There were also pathological changes noted in the muscle ECM of mdx/mTR mice. In support of a fibrotic response, the muscle fibroblast markers PDGFRa and FSP1 (S100a4) were upregulated. There was an induction with nearly every gene that encodes an ECM-related protein, although there was some divergence between RNAseq and protein values (Figure 3D). Numerous proteoglycans such as biglycan (Bgn), decorin (Dcn), lumican (Lum), osteopontin (Spp1), tenascin C (Tnc), tenascin X (Tnxb), versican (Vcan), vinculin (Vcl), and vimentin (Vim) were upregulated at the transcript or protein level (Figure 3D). These observations are consistent with previous findings in 3 month old mdx and 6 month old mdx^4cv^ animals, which are thought to have a more severe muscle phenotype from the original strain of mdx mice (32, 33). However, type Iα1 and type Iα2 collagen had a lower abundance in mdx/mTR muscles, while type VIα1 collagen protein was not different despite having an elevation in transcript (Figure 3D). In 6 month old mdx^4cv^ animals, an elevation in type Iα1 and type VIα1 collagen protein was observed (33). It is possible that the differences between these results in collagen changes are due to an earlier degenerative stage of the mdx/mTR mice, prior to the onset of more severe fibrosis. Indeed, as fibrillar collagens play a role in longitudinal force transmission, and many proteoglycans assist in organizing the extracellular space that is linked with the dystrophin associated glycoprotein complexes (5), it is possible that the induction in proteoglycans occurs due to the repeated cycles of fiber damage and remodeling. Additionally, an elevation in the matrix metalloproteinases MMP-2, −3, and −14 was also observed, and these may be degrading fibrillar and basal lamina collagens (Figure 3D). This is further supported by predicted activation of MMPs from pathway analysis data (Table 1).

Overall, based on the observations in this study and previous reports, the pathological changes to muscle fibers and the ECM in mdx/mTR mice do not appear to disrupt the intrinsic force generating capacity of muscle fibers. However, these changes likely disrupt lateral force transmission throughout the muscle ECM in mdx/mTR mice. The 3.5-fold lower specific force of muscles of mdx/mTR (39) therefore likely occurs due to disruptions in lateral force transmission between fibers and through the ECM rather than deficits in myofibril force production. While the mdx/mTR mouse displays greater pathological changes than the traditional mdx model (39), myofibril structure and the ability of myofibrils to generate force is maintained in mdx/mTR mice.

### Metabolomics and Lipidomics

To determine whether changes in metabolic genes or proteins led to changes in actual metabolite levels, we performed shotgun metabolomics and lipidomics. Much of the previous work that has explored metabolic alterations that occur in DMD animal models has focused on cardiac muscle (13), but some studies have been performed in skeletal muscle tissue. Limited studies in human subjects have also demonstrated reduced succinate and branched chain amino acids, increased glucose, triglycerides and cholesterol, and no difference in creatine/phosphocreatine or phospholipids in muscle biopsies in patients with DMD compared to controls (42, 43). The muscles from golden retriever canines with a form of muscular dystrophy demonstrate reduced Krebs cycle intermediates compared to wild type animals (1) and similar reductions in oxidative capacity were also observed in the muscles of mdx mice (11). In the current study, we detected 62 metabolites in an untargeted mass spectroscopy platform. PC analysis demonstrated moderate overlap between the WT and mdx/mTR metabolome (Figure 6A). We observed higher levels of metabolites involved with glycolysis in mdx/mTR mice, such as 6-phosphoglycerate (6PG), fructose-6-phosphate and glucose-6-phosphate (F6P+G6P), fructose bisphosphate (FBP), phosphorylated hexoses (hexose-P), and phosphoenolpyruvate (PEP), while the branched chain amino acid valine was not different (Figure 6A-B). Consistent with these changes in glycolytic metabolites, enzymes involved in glycolysis such as pyruvate kinase (Pkm) were upregulated, although no difference in aldolase A (Aldoa) was observed, and phosphoglycerate kinase 1 (Pgk1) and phosphorylase kinase (Phkb) were downregulated at the transcript level but protein abundance was not different (Figure 3E). Pathway enrichment analysis identified an induction of glycolysis based on proteomics data, but not from transcriptomics assessments (Table 1).

**Figure 6.**
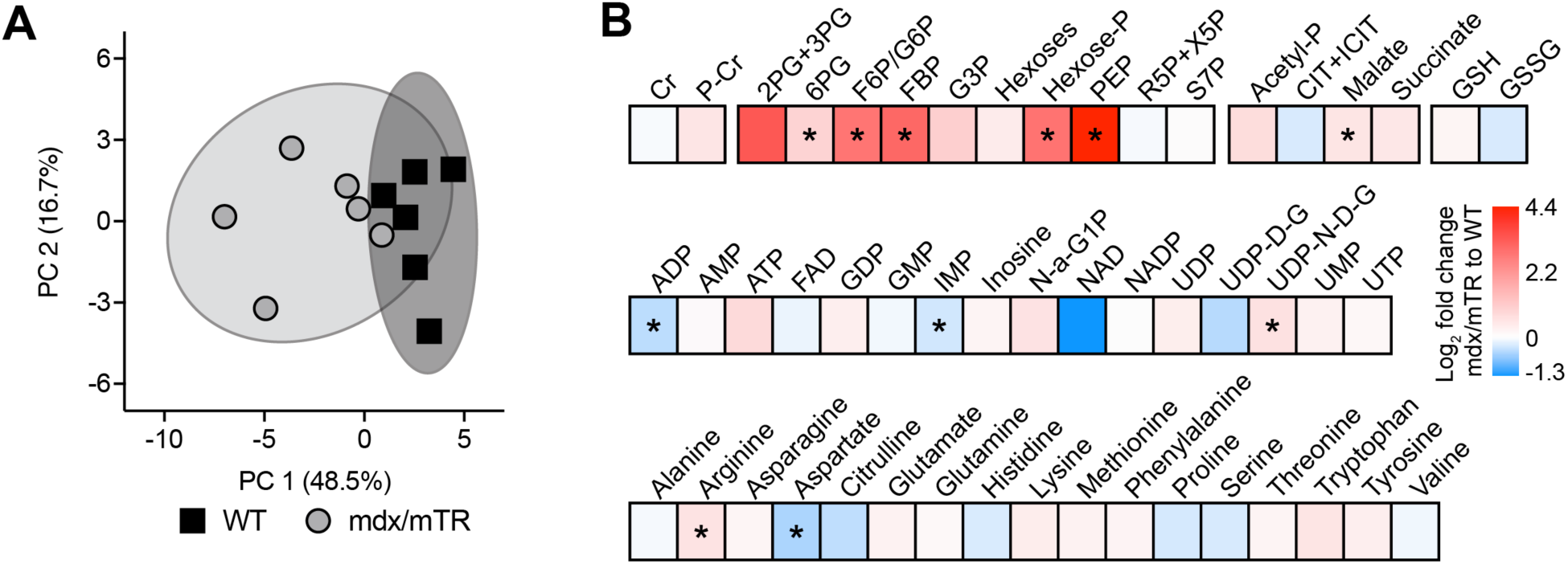
Metabolomics. (A) Principal component (PC) analysis of metabolomics data. (B) Heatmap demonstrating the log_2_ fold change in selected metabolites involved in the creatine phosphate shuttle, glycolysis, Krebs cycle, glutathione metabolism, nucleotide and nucleoside metabolism, as well as free amino acids: creatine (Cr), phospho-creatine (P-Cr), 2-phosphoglycerate and 3-phosphoglycerate (2PG+3PG), 6-phosphoglycerate (6PG), fructose-6-phosphate and glucose-6-phosphate (F6P+G6P), fructose bisphosphate (FBP), glyceraldehyde 3-phosphate (G3P), hexose-phosphates (Hexose-P), phosphoenolpyruvate (PEP), ribulose 5-phosphate and xylulose 5-phosphate (R5P+X5P), sedoheptulose 7-phosphate (S7P), acetyl-phosphate (Acetyl-P), citrate and isocitrate (CIT+ICIT), reduced glutathione (GSH), oxidized glutathione (GSSG), adenosine diphosphate (ADP), adenosine monophosphate (AMP), adenosine triphosphate (ATP), flavin adenine dinucleotide (FAD), guanosine diphosphate (GDP), guanosine monophosphate (GMP), inosine monophosphate (IMP), N-Acetyl-glucosamine-1-phosphate (N-A-G1P), nicotinamide adenine dinucleotide (NAD), nicotinamide adenine dinucleotide phosphate (NADP), uridine diphosphate (UDP), uridine diphosphate-D-glucose (UDP-D-G), uridine diphosphate-N-acetyl-D-glucosamine (UDP-N-D-G), uridine monophosphate (UMP), uridine triphosphate (UTP). Differences between groups assessed using t-tests; *, significantly different (P<0.05) from WT. N=6 WT and N=6 mdx/mTR mice.

For Krebs cycle intermediates, no differences were observed between groups for acetyl-phosphate (Acetyl-P), citrate and isocitrate (CIT+ICIT), or succinate, although malate was elevated (Figure 6B). Aconitase (Aco2) was not different at the transcript or protein level, while citrate synthase (Cs) and succinate dehydrogenase A (Sdha) transcripts were downregulated but had no differences in protein abundance, and myoglobin (Mb) RNA and protein were reduced in mdx/mTR muscles (Figure 3E). Additionally, neuronal nitric oxide synthase (nNOS or Nos1) is an enzyme located at the dystrophin glycoprotein complex that converts the amino acid L-arginine to citrulline, and is known to be disrupted in patients with DMD (5). We observed a downregulation in nNOS gene expression (Figure 3E) along with an accumulation in L-arginine (Figure 6B), consistent with a functional deficit in nNOS activity in mdx/mTR mice. Overall, while markers of glycolysis were enriched in mdx/mTR mice, differences in the Krebs cycle are less clear, and little changes in free amino acids were observed.

Finally, we analyzed the lipid content of muscles. PC analysis demonstrated a general dissimilarity between the lipidome of WT and mdx/mTR mice (Figure 7A). Of the 452 lipid species detected, there were 11 (2.4%) that were at least 1.5-fold elevated and significantly different in mdx/mTR muscles compared to WT, and 20 (4.4%) that were 1.5 fold lower and significantly different (Figure 7B). When analyzing lipid species by class instead of metabolite, ceramides (Cer), diglycerides (DG) and free fatty acids (FFA) were reduced in mdx/mTR muscles, with an elevation in phosphatidylglycerols (PG) and sphingomyelins (SM) (Figure 7C). These findings differ from a previous report of 3 month old mdx mice, which displayed an elevation of PC and PE, although both mdx and mdx/mTR mice did not have differences in PS abundance (35). Most of the enzymes involved in lipid uptake and storage were reduced at the RNA and protein level, such as fatty acid translocase (Cd36), acyl co-A synthetases (Acsl1 and Acss1), diacylglycerol o-acyltransferase 2 (Dgat2), fatty acid binding protein 3 (Fabp3), hormone sensitive lipase (Lipe), and perilipin 5 (Plin5). Additionally, enzymes involved in fatty acid uptake into mitochondria, such as carnitine acylcarnitine translocase (Slc25a20), and carnitine palmitoyltransferase (Cpt1b) were downregulated. For genes involved in β-oxidation of fatty acyl Co-As within the mitochondria, the medium and very long-chain acyl Co-A dehydrogenases (Acadm and Acadvl) were downregulated. Enzymes that produce protaglandins and leukotrienes, such as leukotriene A4 hydrolase (Lta4h), and prostaglandin E synthase 2 and 3 (Ptges2 and Ptges3) were downregulated, although no difference in protein abundance was detected. Similar to the Krebs pathway, there is not a clear pattern for overall deficiencies in lipid oxidative capacity when combining the transcriptome, proteome and lipidomics data, which is further supported by pathway enrichment analysis (Table 1).

**Figure 7.**
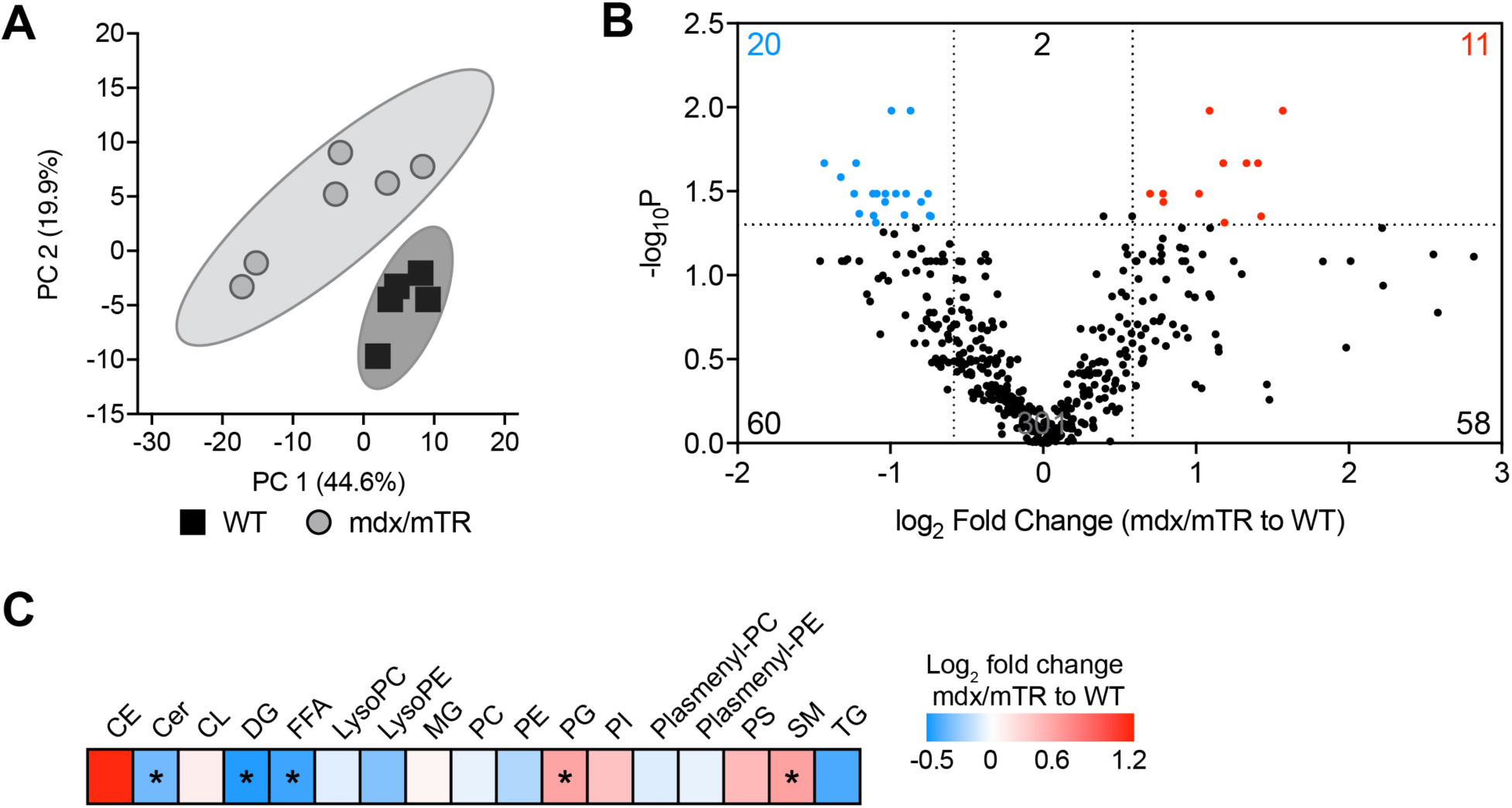
Lipidomics. (A) Principal component (PC) analysis of lipidomics data. (B) Volcano plot demonstrating fold-change and FDR-corrected P-values of all measured lipid species. Lipid species with a > 1.5-fold increase in mdx/mTR mice (log_2_ fold change > 0.585) and P value < 0.05 (-log_10_P > 1.301) are shown in red. Lipid species with a > 1.5-fold decrease in mdx/mTR mice (log_2_ fold change < −0.585) and P value < 0.05 (-log_10_P > 1.301) are shown in blue. (C) Heatmap demonstrating the log_2_ fold change in classes of lipids: Cholesterol ester (CE), ceramides (Cer), cardiolipins (CL), diglycerides (DG), free fatty acids (FFA), lysophosphatidylcholines (LysoPC), lysophosphatidylethanolamines (LysoPE), monoglycerides (MG), phosphatidylcholines (PC), phosphatidylethanolamines (PE), phosphatidylglycerols (PG), phosphatidylinositols (PI), plasmenyl phosphatidylcholines (Plasmenyl-PC), plasmenyl phosphatidylethanolamines (Plasmenyl PE), phosphatidylserines (PS), sphingomyelins (SM), and triglycerides (TG). Differences between groups assessed using t-tests; *, significantly different (P<0.05) from WT. N=5 WT and N=6 mdx/mTR mice.

### Limitations

There are several limitations to this work. We only evaluated mice at a single time point, chosen to be informative about the disease progression, but mdx/mTR mice are known to continue to develop pathological changes as they age. It is possible that extending the analysis to a later time point would demonstrate reductions in muscle fiber force production and additional changes to the transcriptome, proteome, and metabolome. Our analysis focused largely on fast fibered muscles, and other than histology and muscle mass, we did not evaluate changes in slow-fibered muscles like the soleus. We also did not evaluate the diaphragm, which is the most severely injured muscle in mouse models of DMD. Changes in the mRNA transcriptome were studied, and we did not evaluate other types of RNAs that play a role in muscle degeneration and regeneration. Comparisons were made between the transcriptome of the TA and the proteome of the plantaris, which have similar functional and structural properties, but nevertheless are distinct muscles. Mitochondrial respiration or the ability to metabolize glucose, fatty acids, and other substrates were not measured in real time, and conclusions about metabolic changes are based on static RNA, protein, and metabolite measures. We also only evaluated males in this study, as DMD is X-linked, however heterozygous females are known to have some muscle pathologies and reduced physical function. We also used permeabilized muscle fibers to specifically evaluate the contractility of myofibrils, but the combined use of intact and permeabilized muscle fibers would provide more information about the nature of the extent of the force deficit in mdx/mTR mice, and the contributions of metabolic pathways to force production. Despite these limitations, we feel that this work makes an important contribution to our understanding of pathological changes that occur in the muscles of mdx/mTR mice, and identifies several area for future exploration, with particular importance for guiding potential exercise intervention and rehabilitation studies.

### Conclusions

In this study we performed a comprehensive evaluation of the muscle fiber contractile properties, and the transcriptome, proteome, metabolome, and lipidome in WT and mdx/mTR mice in an integrative manner. We found that changes in the proteome only agreed with the transcriptome slightly more than half of the time, and pathway enrichment analysis often diverged between RNAseq and proteomics data. These findings highlight the importance of evaluating changes in the proteome along with the transcriptome. We also provide additional support to the notion that dystrophin is not involved in the longitudinal transfer of force within muscle fibers, and despite extensive inflammation, myofibril function is maintained in mdx/mTR mice. Four months of age appears to be a time in the lifespan of mdx/mTR mice in which fibrotic changes to the ECM are limited, and important metabolic changes were also noted at this time point. The findings of elevated glycolytic metabolism in mdx/mTR mice in this study may also contribute basic science evidence to the emerging field of exercise prescription in patients with DMD (26). Since lengthening contractions are associated with muscle damage, rehabilitation exercises are generally limited to isometric and shortening contractions (21). As low intensity, shortening contractions are associated with causing aerobic adaptations to muscles, and increased aerobic capacity is correlated with overall improved metabolic health and reductions in disease burden (19), low intensity exercise interventions may help patients with DMD to improve their quality of life.

## Supporting information

Supplemental Table S1 - RNAseq

Supplemental Table S2 - Proteomics

Supplemental Table S3 - Metabolomics

Supplemental Table S4 - Lipidomics

Supplemental Table S5 - Primers

Supplemental Table S6 - IPA

## Acknowledgements

The authors would like to acknowledge technical contributions from Mr. Maxwell Konnaris and Mr. Patrick Zager at the Hospital for Special Surgery, and Dr. Deborah Simpson at the University of Liverpool. This work was supported by NIH grants R01-AR063649 and U24-DK097153.

## Author Contributions

DVP, YAK, EC, BM, and CLM designed the study; DVP and CLM wrote the manuscript; DVP, YAK, DCS, LRE, JTD, NPD, ANP, BM, and CLM performed experiments; DVP, YAK, EC, BM, and CLM performed data analysis. All authors reviewed the manuscript.

