## Supplemental Table S5 - Primers for "Multiomics Analysis of the mdx/mTR Mouse Model of Duchenne Muscular Dystrophy"

***Table S5. qPCR Primers.*** Sequences of primers used for qPCR.

| Symbol | Description | GenBank ID | Forward Primer (5' to 3') | Reverse Primer (5' to 3') | Size (bp) |
|---|---|---|---|---|---|
| *B2m* | Beta 2 microglobulin | NM_009735.3 | ATGGGAAGCCGAACATACTG | CAGTCTCAGTGGGGGTGAAT | 177 |
| *Col1a1* | Collagen, type I, alpha 1 | NM_007742.3 | GAGAGGTGAACAAGGTCCCG | AAACCTCTCTCGCCTCTTGC | 153 |
| *Col5a1* | Collagen, type V, alpha 1 | NM_015734.2 | GGAGAGCCACGTGTTCTGTAG | GAGGGAATGAGGCATGGCAG | 135 |
| *Dgat2* | Diacylglycerol O-acyltransferase 2 | NM_026384.3 | ACTGGAACACGCCCAAGAAA | GTAGTCTCGGAAGTAGCGCC | 80 |
| *Fbxo32* | Atrogin-1, F-box protein 32 | NM_026346.3 | GCCCTCCACACTAGTTGACC | GACGGATTGACAGCCAGGAA | 119 |
| *Mmp14* | Matrix metallopeptidase 14 | NM_008608.3 | AGGCCAATGTTCGGAGGAAG | GTGGCACTCTCCCATACTCG | 154 |
| *Mstn* | Myostatin | NM_010834.3 | TCACGCTACCACGGAAACAA | GCAACATTTGGGCTTGCCAT | 82 |
| *Myh8* | Myosin, heavy polypeptide 8, skeletal muscle, perinatal | NM_177369.3 | CGGGAGGTTCACACCAAAAT | CCCTCCTGTGCTTTCCTTCAG | 88 |
| *Tmem8c* | Myomaker, Transmembrane protein 8C | NM_025376.3 | GCCTTTACCACCTTCTCCCCA | GCCTCCATGTAGAAACGCCTC | 144 |
